# Generation of antigen-specific memory CD4 T cells by heterologous immunization enhances the magnitude of the germinal center response upon influenza infection

**DOI:** 10.1101/2023.08.29.555253

**Authors:** Linda M. Sircy, Andrew G. Ramstead, Hemant Joshi, Andrew Baessler, Ignacio Mena, Adolfo García-Sastre, Matthew A. Williams, J. Scott Hale

## Abstract

Current influenza vaccine strategies have yet to overcome significant obstacles, including rapid antigenic drift of seasonal influenza viruses, in generating efficacious long-term humoral immunity. Due to the necessity of germinal center formation in generating long-lived high affinity antibodies, the germinal center has increasingly become a target for the development of novel or improvement of less-efficacious vaccines. However, there remains a major gap in current influenza research to effectively target T follicular helper cells during vaccination to alter the germinal center reaction. In this study, we used a heterologous infection or immunization priming strategy to seed an antigen-specific memory CD4+ T cell pool prior to influenza infection in mice to evaluate the effect of recalled memory T follicular helper cells in increased help to influenza-specific primary B cells and enhanced generation of neutralizing antibodies. We found that heterologous priming with intranasal infection with acute lymphocytic choriomeningitis virus (LCMV) or intramuscular immunization with adjuvanted recombinant LCMV glycoprotein induced increased antigen-specific effector CD4+ T and B cellular responses following infection with a recombinant influenza strain that expresses LCMV glycoprotein. Heterologously primed mice had increased expansion of secondary Th1 and Tfh cell subsets, including increased CD4+ T_RM_ cells in the lung. However, the early enhancement of the germinal center cellular response following influenza infection did not impact influenza-specific antibody generation or B cell repertoires compared to primary influenza infection. Overall, our study suggests that while heterologous infection/immunization priming of CD4+ T cells is able to enhance the early germinal center reaction, further studies to understand how to target the germinal center and CD4+ T cells specifically to increase long-lived antiviral humoral immunity are needed.

**Author Summary:** T follicular helper (Tfh) cells are specialized CD4+ T cells that provide help to B cells and are required to form germinal centers within secondary lymphoid organs during an immune response. Germinal centers are necessary for generating high affinity virus-specific antibodies necessary to clear influenza infections, though current vaccines fail to generate long-lived antibodies that universally recognize different influenza strains. We used a “heterologous priming” strategy in mice using a non-influenza viral infection or viral protein subunit vaccination to form memory CD4+ Tfh cells (in previously naïve mice) that can be rapidly recalled into secondary Tfh cells following influenza infection and ideally enhance the germinal center reaction and formation of high affinity antibodies to influenza better than primary Tfh cells. Our study showed that heterologous priming induced an increase in both CD4+ T and B cells early following influenza infection, suggesting we could successfully target enhancement of the germinal center. Despite the enhancement of the early germinal center cellular response, we did not see an increase in influenza-specific antiviral antibodies. Thus, while Tfh cells are critical for the generation of high affinity antibodies, other strategies to target expansion of Tfh cells during influenza vaccination will need to be developed.

## Introduction

Despite the availability of a vaccine, seasonal influenza infection continues to be a significant burden on the healthcare system in the United States, causing acute respiratory illness and leading to exacerbation of severe health conditions, hospitalization, and mortality (1–3). While current seasonal influenza vaccines prevent millions of influenza-related illness cases each year (2), they fail to induce long-term immunity due to waning neutralizing antibody titers within a year post-vaccination (4–9). (4–9). Additionally, seasonal influenza vaccines fail to induce cross-reactive neutralizing antibodies to the immunodominant globular head of the hemagglutinin (HA) surface glycoprotein due to rapid antigenic drift (10, 11). Currently, one of the major priorities for development and improvement of vaccine strategies is to generate broadly neutralizing antibody responses (12–15), though mechanisms driving the generation of these antibodies after viral infection or vaccination are not well understood.

Formation of the germinal center (GC) during an immune response is necessary for generating long-lived humoral immunity, making it an important target for development of novel vaccines or the improvement of lower efficacy vaccines, including the seasonal influenza vaccine. The GC is where the critical processes for generating long-lived humoral immunity occur, including somatic hypermutation, selection for high affinity antibodies, class switch recombination, and generation of memory B cells and long-lived plasma cells (16, 17). T follicular helper (Tfh) cells are the primary CD4+ T cell subset that helps B cells promote the GC reaction (16–21), and are required to produce long-lived humoral immune responses. Tfh cells are mainly distinguished by expression of the B cell follicle homing receptor CXCR5 (22–25) and the transcriptional repressor Bcl6, which is required for Tfh cell differentiation (26–28). While natural influenza infection and a number of novel vaccine strategies have been shown to induce increased protection and broadly neutralizing antibodies (10, 15, 29–39), the mechanisms or involvement of CD4+ T cells in GC reaction to generate those broadly neutralizing antibodies were not described.

Previous studies have shown that increased circulating memory Tfh cells in HIV-infected patients and highly functional GC Tfh cells in HIV-immunized rhesus macaques correlate with enhanced production of broadly neutralizing antibodies (40–42). In influenza-related studies, adjuvanted inactivated influenza vaccine was shown to increase GC responses and enhance cross-reactivity and long-term detectability of HA-specific antibodies (43). In addition, vaccination with influenza HA-ferritin nanoparticles showed a positive correlation between increased ferritin-specific CD4+ GC Tfh cells and increased HA-specific GC B cells and antibody secreting cells (44). Overall, these studies suggest that current influenza vaccine strategies are ill-equipped at generating universal broadly neutralizing antibodies, possibly in part to mechanisms such as epitope masking by pre-existing antibodies (11, 45–47). Given that Tfh cells have been shown to be the limiting cell subset in the GC reaction (48–50), as well as Tfh cell magnitude correlating with formation of broadly neutralizing antibodies (40–43), we proposed that by directly manipulating Tfh cell magnitude in mice via heterologous infection/immunization priming of CD4+ T cells we would enhance the germinal center reaction and its products compared to primary influenza infection alone.

In this study, we generated antigen-specific CD4+ Tfh memory cells using heterologous priming with either intranasal (i.n.) infection with acute lymphocytic choriomeningitis virus (LCMV) or intramuscular (i.m.) immunization with adjuvanted recombinant glycoprotein from LCMV (rGP) prior to intranasal infection with a recombinant mouse-adapted PR8 strain engineered to carry the CD4-immunodominant LCMVgp61-80 epitope (PR8-HA-GP_61-80_). We then assessed the GC response and antibodies following influenza challenge. We found that heterologous influenza rechallenge resulted in significant increases in the numbers of polyclonal effector antigen-specific CXCR5– Th1 cells in both rGP- and LCMV-primed mice, as well as CXCR5+BCL6+ GC Tfh cells in LCMV-primed mice compared to primary influenza infection. In addition, we analyzed lung-resident CD4+ T cells following heterologous influenza rechallenge and found a significant bias in resident Th1-like cells in LCMV-primed mice and in resident Tfh-like cells in rGP-primed mice, as well as a significant increase in the long-term CD4+ T resident memory pool compared to primary influenza infection. While heterologous infection/immunization priming of CD4+ T cells was able to enhance the early GC cellular response following influenza challenge, we did not see corresponding increases in generating long-term HA-specific antibodies or antibody-secreting cells. Along with previous studies showing the importance of CD4+ Tfh cells in GC and formation of high affinity humoral immunity, our findings suggest that targeting the expansion of memory CD4+ T cells to enhance the primary GC B cell response and tissue-resident memory population is possible and could be a promising avenue to the expansion of memory generation in next generation influenza vaccines.

## Materials and methods

### Viral infections and protein immunizations

C57BL/6J mice (Jackson Laboratory, Bell Harbor, ME) were infected with either 30 μl of 500 TCID_50_ mouse-adapted PR8-HA-GP_61-80_ or 2×10^5^ PFU of LCMV Armstrong by intranasal inoculation or immunized by intramuscular (quadriceps) injection with 2 μg LCMV recombinant glycoprotein (rGP) with addition of Addavax (InvivoGen) adjuvant at a 1:1 ratio. PR8-HA-GP_61-80_ recombinant virus strain is the H1N1 PR8 strain with the CD4-immunodominant LCMVgp61-80 epitope inserted into the HA region and was kindly provided by Dr. Florian Krammer (Icahn School of Medicine at Mount Sinai). 293A cells that express recombinant glycoprotein (from LCMV) were kindly provided by Dr. Carl Davis (Emory University), and recombinant glycoprotein was purified from supernatants as described previously (51). For B cell reactivation experiments, mice were immunized by intraperitoneal injection with 10 μg recombinant HA (H1 subtype) protein from PR8 (H1N1) virus without adjuvant. For intranasal infections, mice were anesthetized with concurrent administration of aerosolized isoflurane and oxygen using a COMPAC^5^ Anesthesia Center (VetEquip). Prior to euthanasia, mice were intravenously injected by retro-orbital injection with 2 μg α-CD45-FITC (30-F11, Tonbo Biosciences) antibody to detect remaining circulating cells in lung samples (52). Animal experiments were conducted in accordance with approved University of Utah IACUC protocols.

### Construction of the recombinant influenza virus PR8-HA-GP_61-80_

To obtain a recombinant influenza virus containing the LCMV epitope GP_61-80_ (GLKGPDIYKGVYQFKSVEFD) inserted in the hemagglutinin (HA) protein, the sequence encoding the GP _61-80_ peptide was introduced in the rescue plasmid pDZ-HA, strain A/Puerto Rico/8/1934 (H1N1) (PR8). The epitope was inserted in-frame at the amino acid position 135, that is highly tolerant to small insertions (53). Next, the recombinant virus was rescued by transfecting cells with 8 plasmids containing the sequences of the viral segments, as previously described (54).

### Tissue processing

Single-cell suspensions of pooled mediastinal lymph nodes or pooled inguinal and lumbar lymph nodes were prepared using 70-μm cell strainers. Single-cell suspensions of spleens were prepared using 70-μm cell strainers and red blood cells lysed by incubation in Ammonium-Chloride-Potassium (ACK) Lysing Buffer (Life Technologies). Single-cell suspensions of lungs were prepared by digestion with 0.25mg/ml Collagenase IV and 15 μg/ml DNase for 1 hour at 37°C, then manually homogenized and red blood cells lysed by incubation in ACK Lysing Buffer and then cells were filtered using 70-μm cell strainers. Cell suspensions were resuspended in RPMI 1640 media supplemented with 5% fetal bovine serum (FBS) prior to FACS staining.

### FACS analysis

Single-cell suspensions of spleens, lungs, and lymph nodes were prepared and up to 2×10^6^ cells were stained in 1X PBS supplemented with 2% fetal bovine serum (FACS buffer) for 15-30 minutes on ice with fluorochrome-conjugated antibodies. Antibodies for FACS included LIVE/DEAD™ Fixable Near-IR Dead Cell Stain, CD4 (RM4-5), CD8 (53-6.7), CD44 (IM7), IFNγ (XMG1.2), TNFα (MP6-XT22), IL-2 (JES6-5H4), PD-1 (29F.1A12), Ly6c (HK1.4), Bcl6 (K112-91), Tbet (4B10), CD19 (eBio1D3 (1D3)), B220 (RA3-6B2), Fas/CD95 (Jo2), GL7 (GL7), IgD (11-26c.2a), CD138 (281–2) (purchased from BD Biosciences, eBiosciences, BioLegend, Vector Laboratories Inc., and Invitrogen). For I-A^b^:gp66-77 tetramer (provided by the National Institutes of Health Tetramer Core) staining, cells were incubated with tetramer in RPMI medium supplemented with 10% FBS for 2 h at 37°C with 5% CO_2_. CXCR5 surface staining was performed using a three-step protocol described in Johnston et al. (2009) (26) using purified rat anti-mouse CXCR5 primary antibody (BD Biosciences, 2G8) in FACS buffer supplemented with 1% bovine serum albumin (Sigma, #A7284) and 2% normal mouse serum (Sigma, #M5905) (CXCR5 staining buffer), a secondary Biotin-SP-conjugated Affinipure F(Ab’)_2_ Goat anti-Rat IgG (Jackson ImmunoResearch) in CXCR5 staining buffer and then with a fluorochrome-conjugated streptavidin in FACS buffer. For transcription factor staining, cells were first stained for surface antigens, followed by permeabilization, fixation and staining using the Foxp3 Permeabilization/Fixation kit and protocol (eBiosciences). Intracellular cytokine staining was done by standard techniques following 5-hour stimulation with Gp_61-80_ peptide and Brefeldin A (GolgiPlug™, BD Biosciences). No peptide controls were treated under the same conditions supplemented with Brefeldin A but without Gp_61-80_ peptide. Cells were then stained for surface antigens, followed by permeabilization, fixation and staining using the Cytofix/Cytoperm kit and protocol (BD Biosciences). For influenza HA-specific B cell staining, recombinant HA protein from A/Puerto Rico/8/1934 (H1N1) virus strain (Immune Technology Corp., #IT-003-0010ΔTMp) was biotinylated with 80-fold molar excess of NHS-PEG4-Biotin solution from the EZ-Link™ NHS-PEG4-Biotin kit (ThermoFisher, #A39259). Excess biotin was removed by buffer exchange of protein into sterile 1X PBS using Zeba™ Spin Desalting Columns, 7K MWCO (ThermoFisher, #89882). Cells were stained on ice for 30min in FACS buffer with 1:100 dilutions of biotin-conjugated-HA and purified rat anti-mouse CD16/CD32 (Mouse BD Fc Block™, Clone 2.4G2, BD Biosciences), then stained on ice for 30min in FACS buffer with 1:1000 dilution of allophycocyanin (APC)-conjugated streptavidin. Cells were analyzed on LSRFortessa™ X-20 and LSRFortessa™ (BD Biosciences) cytometers. FACS data were analyzed using FlowJo v10 software (Tree Star).

### Hemagglutination inhibition assay (HAI)

Serum was separated from whole blood by centrifugation at 10,000xg for 30min at 4°C. HAI to determine neutralizing antibody titers was performed by incubating 25 μL of two-fold serially diluted serum with 25 μL of 4 agglutinating doses (4AD) of WT PR8 (H1N1) virus strain for 30min at room temperature (RT) prior to addition of 50μL of 1% chicken red blood cells (cRBCs) (Lampire Biological Laboratories) in 1X PBS. Plates were gently agitated to mix and then incubated for 30min at RT. HAI titers were determined as the reciprocal dilution of the final well which contained non-agglutinated cRBCs. Naïve mouse serum was used as a negative control. Mice with titers of <1:10 were not included in final analyses.

### Enzyme-linked immunosorbent assay (ELISA)

ELISA to determine HA-specific IgG antibody titers was performed by coating MaxiSorp Clear Flat-Bottom Immuno Nonsterile 96-Well Plates (ThermoFisher) with 1 μg/mL of recombinant HA protein from A/Puerto Rico/8/1934 (H1N1) virus strain (Immune Technology Corp., #IT-003-0010ΔTMp) overnight at 4°C. Plates were blocked for 90min at RT with a solution of 1X PBS with 0.05% Tween® 20 and 10% fetal bovine serum (blocking solution). Plates were incubated with three-fold serially diluted serum in technical duplicates for 90min at RT. Plates were then incubated for 90min at RT with goat anti-mouse IgG conjugated to horseradish peroxidase (HRP) (Southern Biotech, #1030-05) at 1:5000 dilution in blocking solution. Plates were washed with 1X PBS with 0.05% Tween® 20 (PBST) after each blocking/incubation step. Plates were then incubated with 100 μL of substrate solution consisting of 4 mg *o*-Phenylenediamine dihydrochloride (OPD, Sigma, #P8787) dissolved in 10 mL filter sterilized citrate buffer (0.05M citric acid anhydrous, 0.1M sodium phosphate dibasic anhydrous (Na2HPO4)) and 33 μL of 3% H_2_O_2_. The reaction was stopped after 10 min with 100 μL of 1M hydrochloric acid and plates were scanned at 490nm using a Biotek Synergy H1 microplate reader. Naïve mouse serum was used as negative controls. OD readings were averaged between technical duplicates for all samples. Titer cutoff value was determined using the OD values of negative controls as described in Frey et al. (1998) (55) using a 95% confidence level. Relative endpoint titers were calculated by nonlinear regression interpolation of a standard curve (Sigmoidal, 4PL, X is concentration) of individual samples using GraphPad Prism version 9.4.1 for macOS and calculating the titer at which each curve crosses the background cutoff value.

### Enzyme-linked immunosorbent spot assay (ELISpot)

Bone marrow was collected from femur and tibia bones and red blood cells lysed by incubation in ACK Lysing Buffer. B cell enrichment of bone marrow cells was performed using the Pan B Cell Isolation Kit (Miltenyi Biotec, #130-095-813). MultiScreen-IP Filter Plates (Sigma, #MAIPS4510) were pre-wet with 15 μL 35% ethanol for 30 sec and washed with 1X PBS. Plates were coated with 2 μg/mL of recombinant HA protein from A/Puerto Rico/8/1934 (H1N1) virus strain (Immune Technology Corp.) overnight at 4°C. Plates were washed with 1X PBS and then blocked for 2hr at RT with RPMI 1640 medium supplemented with 10% fetal bovine serum, 1% Penicillin-Streptomycin, 2 mM L-glutamine (complete culture medium). Plates were washed with 1X PBS and then enriched B cells in complete culture medium were added to plates at two-fold serial dilutions and in technical duplicates for each sample at maximum 4×10^6^ cells/well and incubated overnight at 37°C at 5% CO_2_. Plates were washed with 1X PBS, then washed with 1X PBS with 0.05% Tween® 20, then incubated with Goat anti-Mouse IgG (H+L) Cross-Adsorbed Secondary Antibody conjugated to HRP (ThermoFisher, #G-21040) at a 1:350 dilution in 1X PBS with 0.05% Tween® 20 and 1% fetal bovine serum overnight at 37°C at 5% CO_2_. Plates were washed with 1X PBS with 0.05% Tween® 20, then washed with 1X PBS, and plates were developed using the AEC Staining Kit (Sigma, #AEC101-1KT). After spot development, plates were washed with water and allowed to dry before counting.

### B cell repertoire sequencing

Single-cell suspensions of splenocytes from individual mice were prepared and B cell enrichment was performed using the Pan B Cell Isolation Kit (Miltenyi Biotec, #130-095-813). Plasmablast cell sorting was performed using a FACSAria (BD Biosciences). Genomic DNA was isolated from sorted plasmablasts using QIAamp DNA Mini Kit (Qiagen), and amplification and sequencing of the *Igh* locus were performed using the immunoSEQ platform (Adaptive Biotechnologies). Data analyses were conducted in the immunoSEQ Analyzer (Adaptive Biotechnologies) and R (56), RStudio (57), and the Immunarch package (58). Data was exported from RStudio using the writexl package (59). Figures were created with the Immunarch package or GraphPad Prism version 9.4.1 for macOS.

### Statistical analysis

All experiments were analyzed using GraphPad Prism version 9.4.1 for macOS. Statistically significant *p* values of <0.05 are indicated and were determined using either a two-tailed unpaired Student’s t test with Welch’s correction or Mann-Whitney U test. Error bars represent Mean±SEM, \**p*≤0.05, \*\**p*≤0.01, \*\*\**p*≤0.001, \*\*\*\**p*≤0.0001.

## Results

### Generation of antigen-specific memory Tfh cells in rGP-immunized and LCMV-infected mice prior to influenza challenge

Our goal was to generate antigen-specific memory CD4+ T cells by heterologous priming with protein immunization or viral infection that could provide help during a primary influenza response. We first evaluated the kinetics and differentiation of polyclonal antigen-specific CD4+ T cells following adjuvanted protein immunization or acute viral infection. C57BL/6J mice were primed by i.m. immunization with 2 μg recombinant LCMV glycoprotein (rGP) in AddaVax adjuvant (GP(1°) group) or by i.n. inoculation with 2×10^5^ PFU acute LCMV-Armstrong (LCMV(1°) group) (**Fig 1A**). At 8, 15, and 39 days post-infection or -immunization (dpi) we analyzed CD4+ T cells in draining lymph nodes – pooled inguinal and lumbar (dLN) following i.m. rGP immunization or mediastinal (medLN) following i.n. LCMV infection – and spleens by staining with the LCMV I-A^b^:gp66-77 MHC class II tetramer. At day 39 post-infection or -immunization, memory I-A^b^:gp66-77 tetramer+ CD4+ T cells were detected in draining lymph nodes in both the GP(1°) and LCMV(1°) groups (**Fig 1B**). In addition, the longitudinal kinetics of I-A^b^:gp66-77 tetramer+ CD4+ T cells analyzed were similar in the draining lymph nodes and spleens of the GP(1°) and LCMV(1°) groups, with peak clonal expansion at 8 dpi and maintenance of the post-contraction memory tetramer+ CD4+ T cell pool detectable at 39 dpi (**Figs S1A-B**). Tetramer+ memory CXCR5+ Tfh cells were similar by frequency and number in the draining lymph nodes of the GP(1°) and LCMV(1°) groups, though the LCMV(1°) group had significantly higher numbers of these cells in the spleen (**Figs 1C-D**).

**Figure 1.**
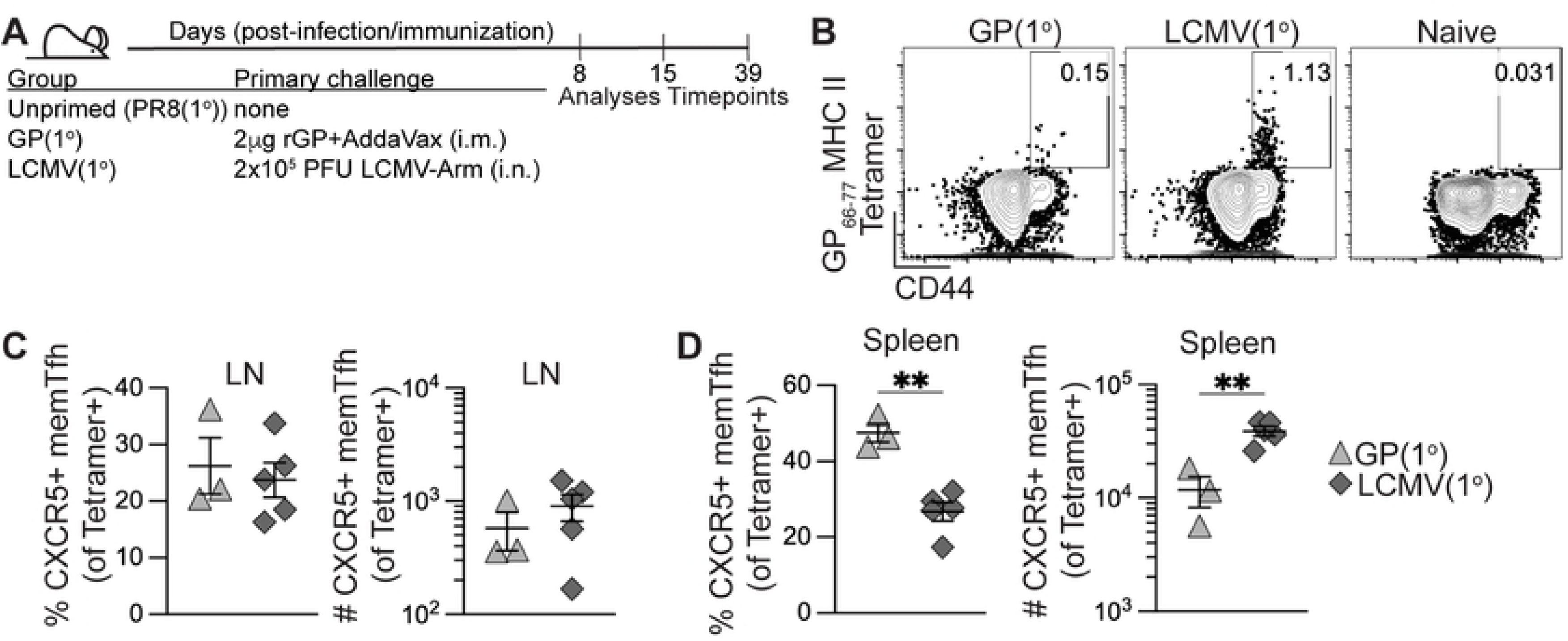
Polyclonal memory CD4+ T follicular helper cell formation following recombinant protein immunization and acute viral infection. C57BL/6J mice were immunized i.m. with 2 μg rGP in AddaVax adjuvant (GP(1°), filled triangle) or infected i.n. with 2×10^5^ PFU of LCMV-Armstrong (LCMV(1°), filled diamond). 8-, 15-, and 39-days postinfection or -immunization, lymphocytes from pooled lumbar and inguinal draining lymph nodes (dLN) (rGP immunization), mediastinal lymph nodes (medLN) (LCMV infection), or spleens were stained with I-A^b^:gp66-77 tetramer. (A) Schematic of experimental design. (B) Representative FACS plots of CD44 and tetramer analysis of total CD4+ T cells in dLN or medLN 39 days postinfection or -immunization. (C) Frequency and number of tetramer+ memory CXCR5+ Tfh cells in dLN or medLN at 39 days postinfection or -immunization. (D) Frequency and number of tetramer+ memory CXCR5+ Tfh cells in spleen at 39 days postinfection or -immunization. *n* ≥ 3 per group per experiment at each timepoint. Data shown are from one independent experiment. Statistically significant *p* values of <0.05 are indicated and were determined using a two-tailed unpaired Student’s t test with Welch’s correction. Error bars represent Mean±SEM, \**p*≤0.05, \*\**p*≤0.01, \*\*\**p*≤0.001, \*\*\*\**p*≤0.0001.

When we analyzed for IFNγ-expressing memory CD4+ T cells in spleen following gp61-80 peptide restimulation and normalized to IFNγ background expression in naïve CD4+ T cells, there was a significantly higher frequency and number of IFNγ-expressing cells in the LCMV(1°) group (**Figs S1C-D**). The lack of IFNγ-expressing cells in the GP(1°) group was expected, as we previously reported that adjuvanted rGP immunization induces expansion of a CXCR5–IFNγ– nonpolarized T helper cell population in lieu of highly IFNγ-expressing Th1 cells as seen following LCMV infection (60). Together, these data show that both adjuvanted rGP immunization and LCMV infection induced I-A^b^:gp66-77 tetramer+ memory CD4+ Tfh and non-Tfh cells.

### Previously generated antigen-specific memory CD4+ T cells induced increased effector Th1 and GC Tfh cells upon influenza infection

We next evaluated if using heterologous priming with adjuvanted rGP immunization or i.n. LCMV infection to generate antigen-specific memory CD4+ T cells would enhance the early effector germinal center response to influenza infection. Using the heterologous priming strategies detailed in Figure 1A, 42 days after priming infection or immunization we infected i.n. with 500 TCID_50_ PR8-HA-GP_61-80_ recombinant influenza virus (GP(1°)PR8(2°) and LCMV(1°)PR8(2°) groups) (**Fig 2A**). For control groups, we used age- and sex-matched naïve mice infected i.n. with PR8-HA-GP_61-80_ to evaluate the primary influenza response (PR8(1°) group) and a separate group was homologously primed with PR8-HA-GP_61-80_ i.n. infection (PR8(1°)PR8(2°) group) to evaluate recalled cellular and antibody responses (**Fig 2A**). 8 days after influenza infection, both GP(1°)PR8(2°) and LCMV(1°)PR8(2°) groups had significantly higher frequencies and numbers of effector tetramer+ CD4+ T cells in medLN than the PR8(1°) group, with LCMV-primed mice having the highest overall (**Figs 2B-C**). Heterologous infection/immunization priming prior to influenza infection induced increased frequencies and numbers of effector tetramer+ CXCR5– TBET+ Th1 cells compared to primary influenza infection alone at 8 dpi (**Figs 2D-E**). While PR8(1°) mice had a significantly higher frequency of CXCR5+PD-1+ Tfh and CXCR5+BCL6+ GC Tfh cells in medLN, the numbers of GC Tfh cells were significantly higher in LCMV-primed mice (**Figs 2D, 2F-G**). Although the numbers of BCL6-expressing CXCR5+ GC Tfh cells were highest in LCMV-primed mice, the amount of BCL6 expression in tetramer+ CXCR5+ cells was significantly lower in these mice (**Figs 2G-H**). In addition, our data showed distinct populations in medLN of CXCR5+LY6C^low^ Tfh cells and CXCR5– LY6C^high^ Th1 cells, although the Th1 cells in the GP(1°)PR8(2°) group were mostly LY6C^low^ (**Fig S2A**).

**Figure 2.**
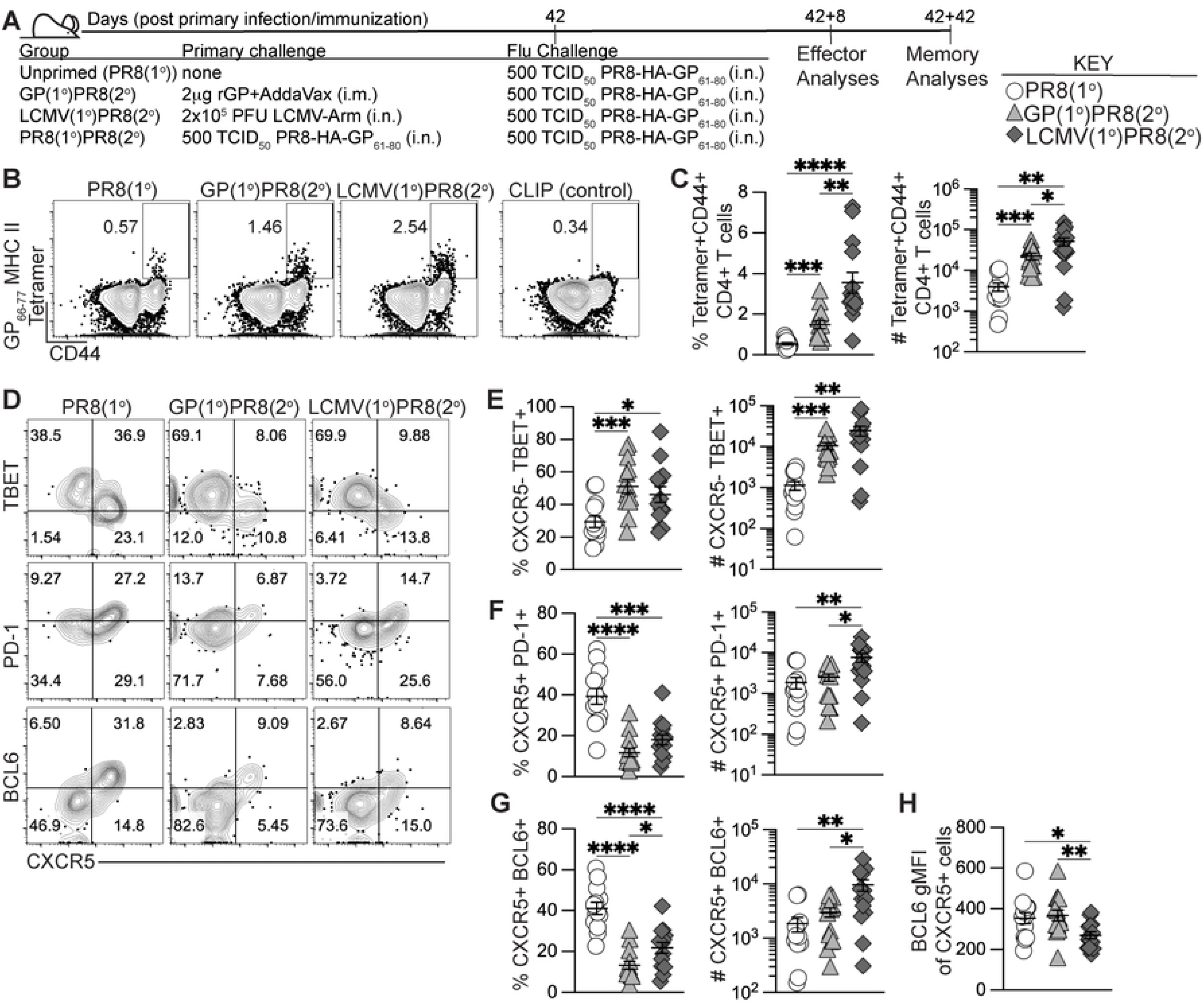
Generation of memory CD4+ T cells by heterologous immunization induced increased effector antigen-specific Th1 and GC Tfh cells following influenza infection. C57BL/6J mice were primed by i.m. immunization with 2 μg rGP in AddaVax (GP(1°)PR8(2°), filled triangle) or by i.n. infection with 2×10^5^ PFU of LCMV-Armstrong (LCMV(1°)PR8(2°), filled diamond). 42 days postinfection or -immunization, primed mice and unprimed age-matched naïve mice (PR8(1°), unfilled circle) were infected i.n. with 500 TCID_50_ of PR8-HA-GP_61-80_ influenza virus. 8 days after influenza infection, lymphocytes from medLN were stained with I-A^b^:gp66-77 tetramer or stained with I-A^b^ human CLIP87-101 as a control. **(A)** Schematic of experimental design. **(B)** Representative FACS plots of CD44 and tetramer analysis of total CD4+ T cells. **(C)** Frequency and number of effector tetramer+CD44+ of total CD4+ T cells. **(D)** Representative FACS plots of CXCR5, TBET, PD-1, and BCL6 analysis of tetramer+CD44+ CD4+ T cells. **(E)** Frequency and number of effector tetramer+ CXCR5–TBET+ T helper 1 cells. **(F)** Frequency and number of effector tetramer+ CXCR5+PD-1+ Tfh cells. **(G)** Frequency and number of effector tetramer+ CXCR5+BCL6+ GC Tfh cells. **(H)** BCL6 geometric mean fluorescence intensity (gMFI) of tetramer+ CXCR5+ cells. *n* ≥ 3 per group per experiment. Data shown are from three independent experiments. Statistically significant *p* values of <0.05 are indicated and were determined using a two-tailed unpaired Student’s t test with Welch’s correction. Error bars represent Mean±SEM, \**p*≤0.05, \*\**p*≤0.01, \*\*\**p*≤0.001, \*\*\*\**p*≤0.0001.

Cytokine analysis of CD4+ T cells in medLN following gp61-80 restimulation revealed that LCMV-primed mice had the highest frequency and number of both IFNγ+ and polyfunctional IFNγ+TNFα+IL-2+ expressing cells (**Figs S2B-D**). The GP(1°)PR8(2°) group also had a significantly higher number of IFNγ+ expressing cells than the PR8(1°) group, further confirming that memory nonpolarized T helper cells can form Th1 cells after secondary activation (**Fig S2C**). We also evaluated differences in tetramer+ Th1 and Tfh cells in the spleen and found that Th1 cells (by both TBET and IFNγ expression) were similar to Th1 cells in medLN, as LCMV-primed mice were significantly higher overall (**Figs S2E-G**). However, our data showed no differences in Tfh cells in the spleen by PD-1 or BCL6 expression (**Figs S2H-I**). Together, our data show that heterologous priming with rGP immunization or LCMV infection had induced enhanced expansion of CD4+ Tfh and Th1 cells compared to primary influenza infection.

### Prior generation of memory Tfh cells promoted increased influenza-specific GC B cells and plasmablasts upon influenza infection

To determine if recalled antigen-specific memory CD4+ T cells in heterologously primed mice could enhance the primary anti-influenza B cell response, we next analyzed GC B cell and plasmablast populations in medLN and spleen. While there were no differences in frequency or number of total CD19+B220+/low B cells in medLN (**Fig S3A**), heterologous infection/immunization priming of CD4+ T cells drove a significant increase in the number of total Fas+GL7+ GC B cells and influenza HA-specific GC B cells (**Figs 3A-D**) at 8 dpi. In addition, while the number of total IgD–CD138+ plasmablasts in medLN were similar between groups, there was a significant increase of HA-specific plasmablasts in both rGP- and LCMV-primed mice compared to PR8(1°) mice (**Figs 3E-H**). Analysis of the splenic B cell response revealed significant increases in numbers of total GC B cells and plasmablasts, as well as HA-specific GC B cells and plasmablasts in the GP(1°)PR8(2°) group compared to the PR8(1°) group (**Figs S3B-F**), despite no differences in splenic tetramer+ GC Tfh cells (**Figs S2H-I**).

**Figure 3.**
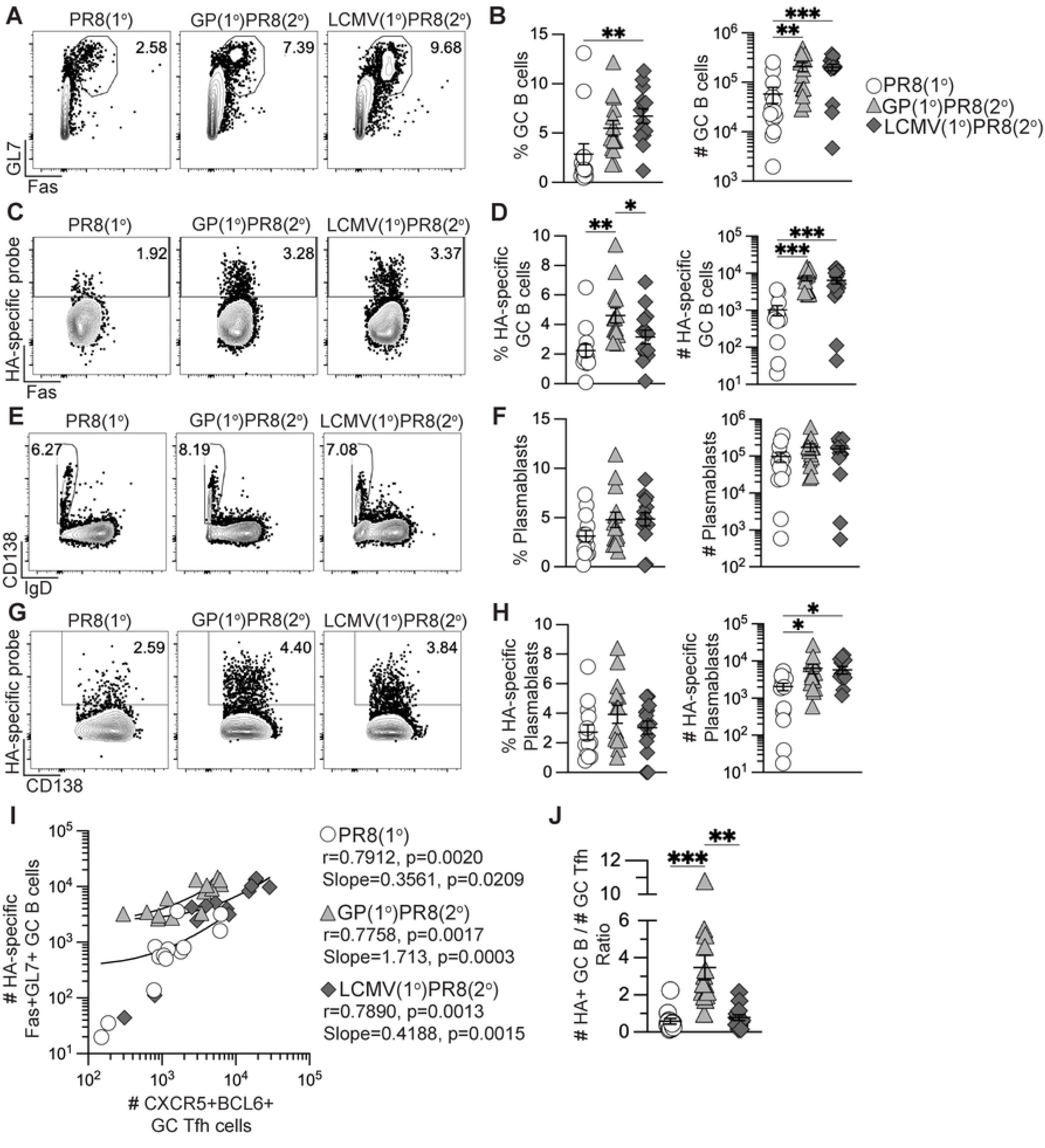
Generation of memory CD4+ T cells by heterologous immunization induced increased influenza-specific B cells following influenza infection. Flow cytometry analysis of B cells from medLN 8 days after PR8-HA-GP_61-80_ influenza virus infection in rGP immunization-primed (GP(1°)PR8(2°), filled triangle), LCMV-primed (LCMV(1°)PR8(2°), filled diamond), or unprimed naïve mice (PR8(1°), unfilled circle). **(A)** Representative FACS plots of Fas and GL7 analysis gated on total CD19+B220+/low cells. **(B)** Frequency and number of Fas+GL7+ GC B cells of total CD19+B220+/low B cells. **(C)** Representative FACS plots of influenza HA-specific GC B cells gated on total Fas+GL7+ GC B cells. **(D)** Frequency and number of HA-specific GC B cells of total Fas+GL7+ GC B cells. **(E)** Representative FACS plots of IgD and CD138 analysis gated on total CD19+B220+/low cells. **(F)** Frequency and number of IgD–CD138+ plasmablasts of total CD19+B220+/low cells. **(G)** Representative FACS plots of influenza HA-specific plasmablasts gated on total IgD–CD138+ plasmablasts. **(H)** Frequency and number of influenza HA-specific plasmablasts of total IgD–CD138+ plasmablasts. **(I)** Correlation analysis of number of tetramer+ CXCR5+BCL6+ GC Tfh cells to number of HA-specific Fas+GL7+ GC B cells. Spearman rank-order correlation values (r) and statistically significant *p* values are shown with linear regression curve fit line slopes and statistically significant *p* values. **(J)** Ratio of number of HA-specific GC B cells to number of tetramer+ CXCR5+BCL6+ GC Tfh cells. *n* ≥ 3 per group per experiment. Data shown are from three independent experiments. Statistically significant *p*values of <0.05 are indicated and were determined using a two-tailed unpaired Student’s t test with Welch’s correction. Error bars represent Mean±SEM, \**p*≤0.05, \*\**p*≤0.01, \*\*\**p*≤0.001, \*\*\*\**p*≤0.0001.

To determine if there was a correlation between the numbers of tetramer+ GC Tfh cells and HA-specific GC B cells in medLN, we performed Spearman correlation analysis and found that in all groups there was a statistically significant positive correlation (**Fig 3I**). In addition, linear regression analysis of GC Tfh and HA-specific GC B cell numbers showed statistically significant positive associations in all groups (**Fig 3I**), consistent with previous studies describing the critical interaction between Tfh and B cells in the germinal center (26, 48, 61–63). We further compared the numbers of HA-specific GC B cells to the number of GC Tfh cells and found that in the GP(1°)PR8(2°) group, there was a significantly higher number of HA-specific GC B cells to every GC Tfh cell (**Fig 3J**). These data suggest that while the number of GC Tfh cells in the GP(1°)PR8(2°) group were similar to the PR8(1°) group, the GC Tfh cells may be of a higher quality as to sustain support for higher numbers of HA-specific GC B cells, as has been previously suggested (41, 44). Together, these data suggest that heterologous priming with adjuvanted rGP immunization or LCMV infection significantly enhanced the early anti-influenza germinal center B cell response following influenza infection compared to primary influenza infection.

### Previously generated memory CD4+ T cells by heterologous immunization did not impact anti-influenza antibody titers

To determine if the early increases in tetramer+ CD4+ T cells and influenza-specific B cells in heterologously primed mice were maintained at memory, we analyzed the cellular response longitudinally in medLN and spleen 42 days after influenza challenge as detailed in Figure 2A. Similar to 8 days after influenza challenge, heterologously primed mice maintained significantly higher frequencies and numbers of I-A^b^:gp66-77 tetramer+ CD4+ T cells in medLN 42 days after influenza challenge compared to the unprimed PR8(1°) and homologously primed PR8(1°)PR8(2°) groups (**Figs 4A and S4A**). Our data also showed significantly greater frequencies and numbers of tetramer+ CXCR5–TBET+ Th1 cells and significantly higher numbers of antigen-specific polyfunctional cytokine-secreting (IFNγ+TNFα+IL-2+) CD4+ T cells in heterologously primed mice in medLN (**Figs 4B-C**). In addition, memory I-A^b^:gp66-77 tetramer+ CD4+ T cells, tetramer+ CXCR5–TBET+ Th1 cells and polyfunctional cytokine secreting CD4+ T cells were detectable in the spleen at significantly higher numbers in heterologously primed mice (**Figs S4B-D**).

**Figure 4.**
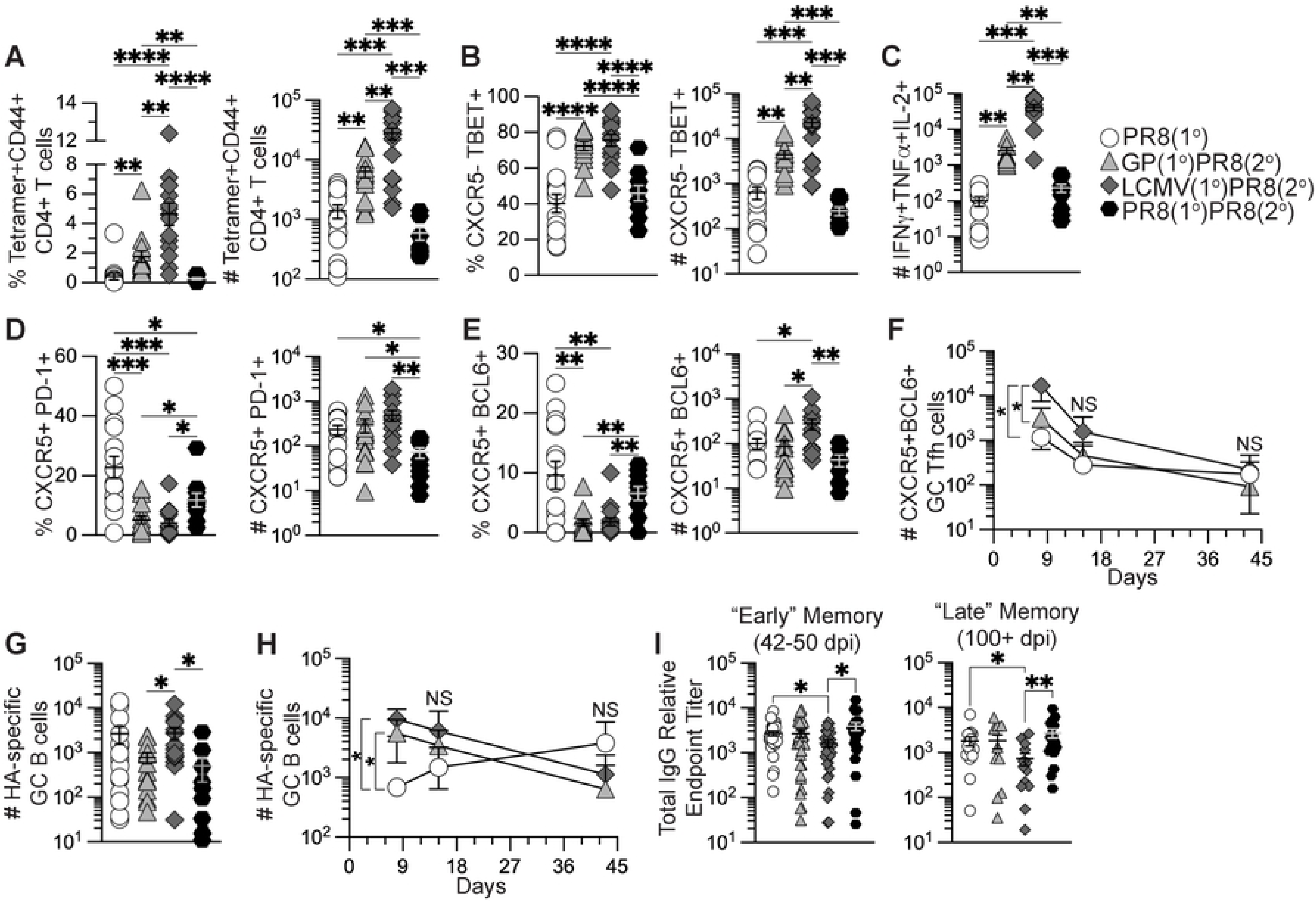
Generation of memory CD4+ T cells by heterologous immunization induced increased memory Th1 cells remaining after influenza infection but did not enhance influenza-specific antibody titers. Flow cytometry analysis of CD4+ T cells and B cells from medLN 42 days after PR8-HA-GP_61-80_ influenza virus infection in heterologously primed mice (GP(1°)PR8(2°), filled triangle, or LCMV(1°)PR8(2°), filled diamond), homologously primed mice (PR8(1°)PR8(2°), filled hexagon), or unprimed naïve mice (PR8(1°), unfilled circle). CD4+ T cells were analyzed by staining with I-A^b^:gp66-77 tetramer and cytokine expression in CD4+ T cells was analyzed following restimulation with gp61-80 peptide. Serum was isolated from whole blood collected from influenza infected mice at 15-16, 42-50, and 100+ days postinfection and analyzed by ELISA. **(A)** Frequency and number of I-A^b^:gp66-77 tetramer+CD44+ of total CD4+ T cells in medLN at 42 days postinfection. **(B)** Frequency and number of I-A^b^:gp66-77 tetramer+ CXCR5–TBET+ T helper 1 cells. **(C)** Number of antigen-specific IFNγ+TNFα+IL-2+ cells. **(D)** Frequency and number of tetramer+ CXCR5+PD-1+ Tfh cells. **(E)** Frequency and number of tetramer+ CXCR5+BCL6+ GC Tfh cells. **(F)** Kinetics of tetramer+ CXCR5-+BCL6+ GC Tfh cells in medLN at 8, 15, and 42 days postinfection. **(G)** Number of HA-specific GC B cells of total Fas+GL7+ GC B cells in medLN at 42 days postinfection. **(H)** Kinetics of HA-specific GC B cells in medLN at 8, 15, and 42 days postinfection. **(I)** Anti-influenza H1 HA-specific IgG antibody titers from serum at 42-50 and 100+ days postinfection by ELISA. *n* ≥ 3 per group per experiment. Kinetics data (panels F and H) shown are from one independent experiment. FACS and serology data shown are from two to three independent experiments. Statistically significant *p* values of <0.05 are indicated and were determined using a two-tailed unpaired Student’s t test with Welch’s correction. Error bars represent Mean±SEM, \**p*≤0.05, \*\**p*≤0.01, \*\*\**p*≤0.001, \*\*\*\**p*≤0.0001. NS=not significant.

As the CD4+ T cell-specific immunodominant LCMVgp61-80 epitope contains a cryptic epitope recognized by CD8+ T cells (64), we analyzed CD8+CD44+ T cells at 8 and 42 days after influenza challenge for IFNγ expression following LCMVgp61-80 peptide restimulation. Our data show that most mice had frequencies of CD44+IFNγ+ cells of CD8+ T cells below background levels as normalized to a no peptide control, and all other mice had less than 1% of CD8+ T cells expressing CD44 and IFNγ (**Fig S4E**). These data suggest that while non-specific secretion of IFNγ by CD8+ T cells was detected, we do not expect these cells significantly influenced the increases in Th1 cells and IFNγ-secreting CD4+ T cells. We next analyzed I-A^b^:gp66-77 tetramer+ CD4+ T cells for CXCR5, PD-1, and BCL6 expression following influenza infection in medLN to determine the kinetics of memory antigen-specific Tfh cells. At 42 days after influenza infection, the PR8(1°) group had maintained a significantly higher frequency of both PD-1- and BCL6-expressing Tfh cells (**Figs 4D-E**) similar to the effector timepoint. While BCL6 expression in memory CXCR5+ Tfh cells is significantly reduced following acute viral clearance compared to effector CXCR5+ GC Tfh cells (65), LCMV(1°)PR8(2°) mice still had significantly higher numbers of BCL6-expressing Tfh cells maintained at memory compared to the other three groups (**Fig 4E**). When we analyzed lymphocytes at 8, 15 and 42 days after influenza infection to evaluate differences in proliferation or contraction kinetics of CXCR5+BCL6+ GC Tfh cells, our data showed no differences in longitudinal kinetics of GC Tfh cells in this experiment and only significantly higher numbers of GC Tfh cells in heterologously primed mice at 8 dpi (**Fig 4F**).

We then analyzed the memory B cell pool in medLN and found only the LCMV(1°)PR8(2°) group had significantly higher numbers of HA-specific GC B cells compared to GP(1°)PR8(2°) and PR8(1°)PR8(2°) mice at 42 dpi (**Fig 4G**). When we analyzed HA-specific GC B cell kinetics at 8, 15, and 42 days after influenza infection, HA-specific GC B cells underwent contraction after peak expansion around 8 dpi in heterologously primed mice (**Fig 4H**). However, in the PR8(1°) group HA-specific GC B cells increased in number after 8 dpi, though numbers were not significantly different from heterologously primed at 15 or 42 dpi in this experiment (**Fig 4H**).

As heterologous infection/immunization priming significantly increased HA-specific GC B cells and plasmablasts 8 days after influenza infection, we analyzed the sera of influenza infected mice to determine HA-specific neutralizing antibody and IgG antibody titers. We found that LCMV infection had a slight but statistically significant adverse impact on HA-specific IgG antibody titers compared to influenza infection alone (**Fig 4I**). In addition, despite heterologous infection/immunization priming inducing increased antigen-specific GC Tfh and GC B cells 8 days after influenza infection, all groups had similar HA-specific neutralizing antibody titers at all timepoints (**Fig S4F**). To determine if the enhanced early germinal center cellular response in heterologously primed mice corresponded to an increase in HA-specific long-lived plasma cells, we analyzed enriched B cells from bone marrow of infected mice for IgG secretion by ELISpot 42 and 105 days after influenza infection. As with our serology data, we found no differences in HA-specific IgG-secreting B cells from infected mice regardless of priming strategy (**Figs S4G-H**). Together these data suggest that while heterologous infection/immunization priming of CD4+ T cells did significantly enhance germinal center CD4+ T and B cell responses early after influenza infection, those effects did not significantly impact long-term germinal center-driven humoral responses compared to primary influenza infection. In addition, our data show that both adjuvanted rGP immunization and LCMV infection significantly enhanced the memory antigen-specific Th1 cell pool after influenza infection, despite differences in the CXCR5– non-Tfh cell populations prior to influenza infection.

### Prior generation of antigen-specific memory CD4+ T cells enhanced early GC responses and long-term lung-resident Th1 cells upon influenza infection

Recent evidence has indicated an important role for CD4+ T resident memory (T_RM_) cells in mediating protection from influenza infection in the lung (66–70). Lung-resident CD4+ T cell responses result in the formation of T_RM_ with either Th1 or Tfh properties that can coordinate localized immune responses (71, 72). Furthermore, lung-specific immune responses are characterized by the induction of localized B cell responses and the formation of long-lived tissue-resident memory B cells that primarily home to the bronchoalveolar lymphoid tissue (BALT), and it is likely that localized antibody responses comprise a key line of defense against influenza infection in the lung (73–75). To determine the impact of heterologous infection/immunization priming of CD4+ T cells on the establishment and boosting of secondary T_RM_ we assessed CD4+ T cell responses in the lung following influenza infection of primed and unprimed mice as previously described in Figure 2A. We employed intravascular anti-CD45 staining to distinguish lung-infiltrating leukocytes from those in circulation (52), combined with CD69 staining to identify T_RM_ (67). Both primary adjuvanted rGP immunization and LCMV i.n. infection induced small numbers of lung-infiltrating memory I-A^b^:gp66-77 tetramer+ CD4+ T cells detected at 39 dpi (**Figs S5A-C**) that were dramatically boosted in frequency and number at 8 days after influenza infection and were significantly higher compared to the PR8(1°) and PR8(1°)PR8(2°) groups (**Figs 5A-B**). The priming strategy utilized also impacted the resulting secondary effector CD4+ T cell subsets. After PR8-HA-GP_61-80_ infection, rGP immunization-induced memory CD4+ T cells preferentially gave rise to FR4+LY6C– Tfh-like secondary effector cells, whereas LCMV-induced memory T cells gave rise to FR4–LY6C+ Th1-like secondary effector cells (**Figs 5C-E**).

**Figure 5.**
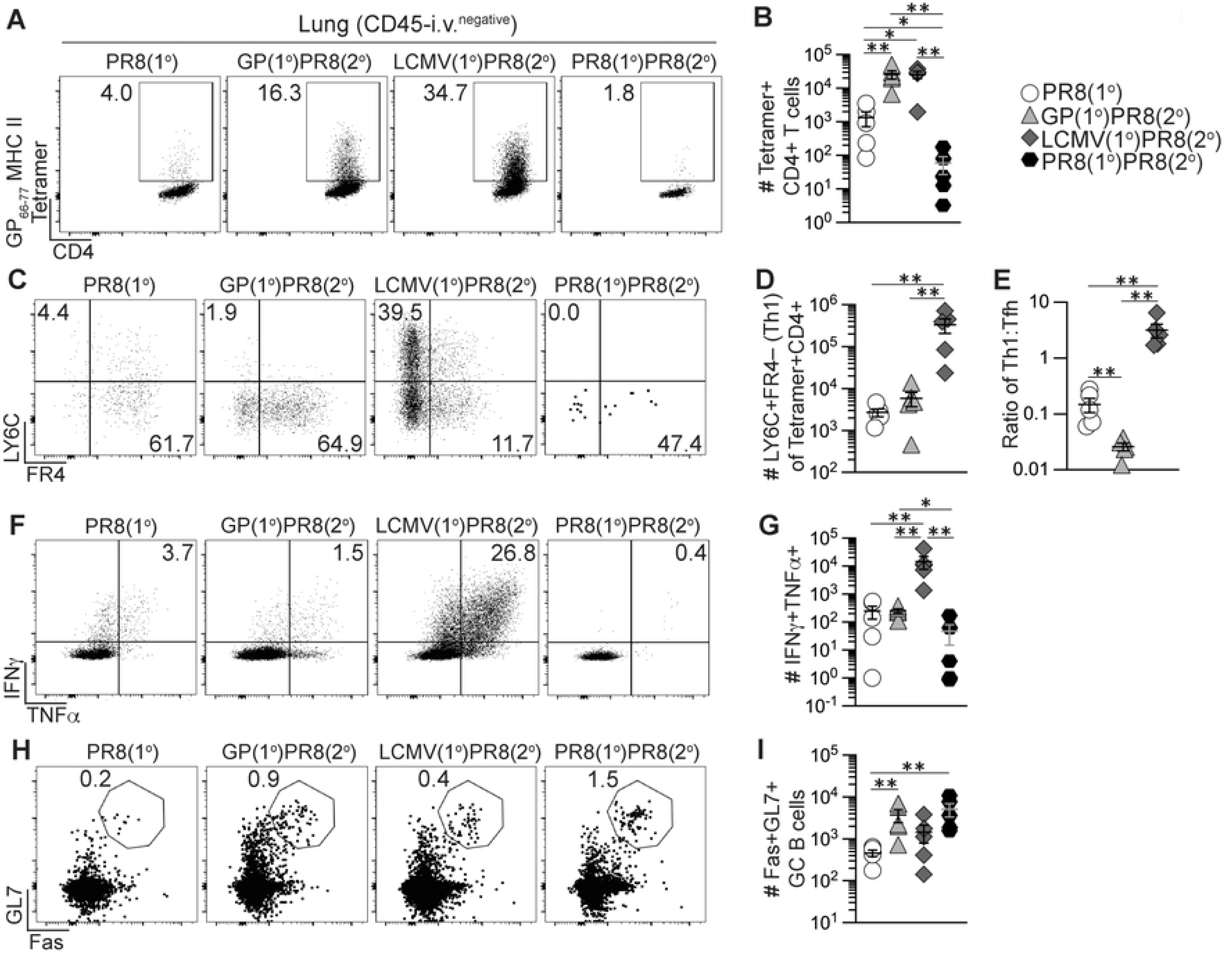
Generation of memory CD4+ T cells by heterologous immunization induced increased GC B cells and Th1 cells in lung early following influenza infection. Flow cytometry analysis of CD45-i.v.^negative^ CD4+ T cells and B cells from lung 8 days after PR8-HA-GP_61-80_ influenza virus infection in heterologously primed mice (GP(1°)PR8(2°), filled triangle, or LCMV(1°)PR8(2°), filled diamond), homologously primed mice (PR8(1°)PR8(2°), filled hexagon) or unprimed naïve mice (PR8(1°), unfilled circle). CD4+ T cells were analyzed by staining with I-A^b^:gp66-77 tetramer and cytokine expression of CD4+ T cells was analyzed following restimulation with gp61-80 peptide. **(A)** Representative FACS plots of I-A^b^:gp66-77 tetramer analysis of total CD4+ T cells. **(B)** Number of I-A^b^:gp66-77 tetramer+ cells of total CD4+ T cells. **(C)** Representative FACS plots of FR4 and LY6C analysis gated on I-A^b^:gp66-77 tetramer+CD4+ T cells. **(D)** Number of tetramer+ LY6C+FR4– (Th1) cells. **(E)** Ratio of the number of tetramer+ LY6C+FR4– (Th1) cells to number of tetramer+ LY6C–FR4+ (Tfh) cells. **(F)** Representative FACS plots of TNFα and IFNγ analysis gated on total CD4+ T cells. **(G)** Number of IFNγ+TNFα+ cells of total CD4+ T cells. **(H)** Representative FACS plots of Fas and GL7 gated on CD19+ B cells. **(I)** Number of Fas+GL7+ GC B cells of total B cells. *n* ≥ 3 per group per experiment at each timepoint. Data shown are from one experiment and are representative of two to three independent experiments. Statistically significant *p* values of <0.05 are indicated and were determined using Mann-Whitney U test. Error bars represent Mean±SEM, \**p*≤0.05, \*\**p*≤0.01, \*\*\**p*≤0.001, \*\*\*\**p*≤0.0001.

Prior to PR8-HA-GP_61-80_ infection, we analyzed lung-infiltrating CD4+ T cells for cytokine expression following *ex vivo* gp61-80 peptide restimulation and found that primary LCMV i.n. infection induced significantly more IFNγ- and TNFα-producing T cells in the lung compared to adjuvanted rGP immunization at 39 dpi (**Figs S5D-E**) similar to our data of CD4+ T cells in the lymph nodes and spleen. When we analyzed effector CD4+ T cells in the lung for cytokine expression 8 days after PR8-HA-GP_61-80_ challenge, the LCMV(1°)PR8(2°) group had the highest expansion of CD4+ T cells producing IFNγ and TNFα (**Figs 5F-G**), despite the presence of similar numbers of total tetramer+ CD4+ T cells to the GP(1°)PR8(2°) group (**Fig 5B**).

We then investigated the impact of heterologous infection/immunization priming of CD4+ T cells on GC B cells in the lung. We found that primary adjuvated rGP immunization and LCMV i.n. infection induced similar numbers of total B cells and GC B cells in the lung prior to PR8-HA-GP_61-80_ challenge detected at 39 dpi (**Figs S5F-I**). However, 8 days after PR8-HA-GP_61-80_ challenge, our data showed that adjuvanted rGP immunization induced significantly more GC B cells (CD19+GL7+Fas+) in the lung, as compared to the PR8(1°) group (**Figs 5H-I**).

We next sought to determine the impact of heterologous infection/immunization priming of CD4+ T cells on the establishment of lung-infiltrating memory CD4+ T cells following influenza infection. As was the case for the secondary effector response in the lung (**Fig 5**), PR8-HA-GP_61-80_ rechallenge of rGP immunization-or LCMV infection-derived memory CD4+ T cells resulted in a large population of tetramer+ secondary memory T cells in the lung at 42 dpi as compared to the PR8(1°) and PR8(1°)PR8(2°) groups (**Figs 6A-B**). Most of these cells expressed CD69, a marker of lung CD4+ T_RM_ following influenza infection (67, 68), resulting in a 50-100-fold increase in CD4+ T_RM_ following heterologous rechallenge of rGP- or LCMV-derived CD4+ memory T cells (**Figs 6C-D**). In addition, the memory CD4+ T cells maintained their primary activation-dependent Th1 and Tfh bias, as the LCMV(1°)PR8(2°) mice had significantly more LY6C+ Th1-like secondary memory T cells 42 days after influenza infection and the GP(1°)PR8(2°) mice had significantly more FR4+ Tfh-like secondary memory T cells (**Figs 6E-G**). Overall, our findings showed that heterologous infection/immunization priming of CD4+ T cells induced large numbers of lung T_RM_ following PR8-HA-GP_61-80_ rechallenge, with a Th1-like or Tfh-like subset distribution that was dependent on the primary immunization or infection challenge.

**Figure 6.**
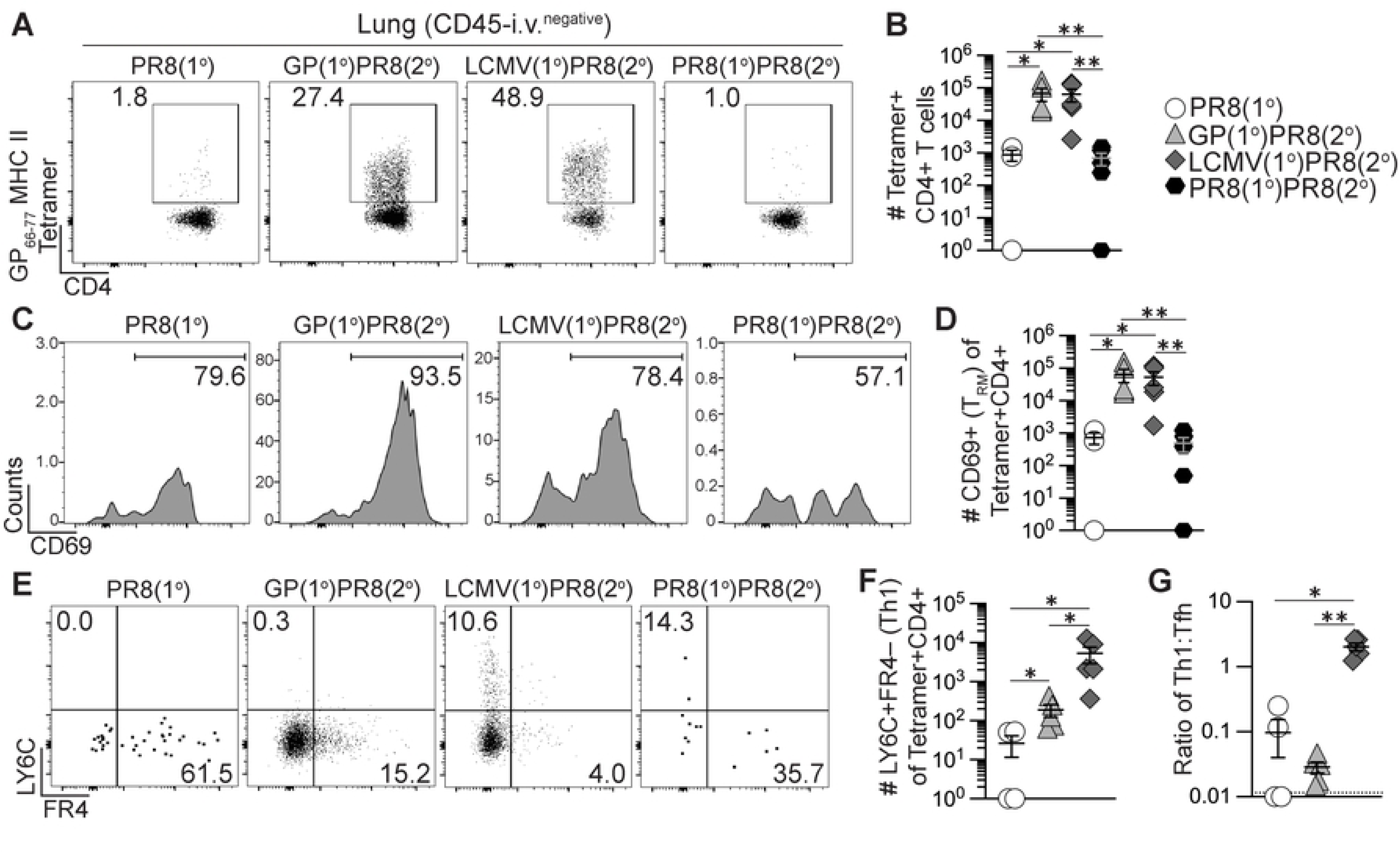
Generation of memory CD4+ T cells by heterologous immunization enhanced long-term memory Th1 and CD4+ T_RM_ cells following influenza infection. Flow cytometry analysis of CD45-i.v.^negative^ CD4+ T cells from lung 42 days after PR8-HA-GP_61-80_ influenza virus infection in heterologously primed mice (GP(1°)PR8(2°), filled triangle, or LCMV(1°)PR8(2°), filled diamond), homologously primed mice (PR8(1°)PR8(2°), filled hexagon) or unprimed naïve mice (PR8(1°), unfilled circle). CD4+ T cells were analyzed by staining with I-A^b^:gp66-77 tetramer. **(A)** Representative FACS plots of CD4 and I-A^b^:gp66-77 tetramer analysis of total CD4+ T cells. **(B)** Number of I-A^b^:gp66-77 tetramer+CD44+ of total CD4+ T cells. **(C)** Representative FACS plots of CD69 analysis gated on I-A^b^:gp66-77 tetramer+CD4+ T cells. **(D)** Number of CD69+ T_RM_ cells of tetramer+CD4+ T cells. **(E)** Representative FACS plots of FR4 and LY6C analysis gated on I-A^b^:gp66-77 tetramer+CD4+ T cells. **(F)** Number of tetramer+ LY6C+FR4– (Th1) cells. **(G)** Ratio of the number of tetramer+ LY6C+FR4– (Th1) cells to number of tetramer+ LY6C– FR4+ (Tfh) cells. *n* ≥ 3 per group per experiment at each timepoint. Data shown are from one experiment and are representative of two to three independent experiments. Statistically significant *p* values of <0.05 are indicated and were determined using Mann-Whitney U test. Error bars represent Mean±SEM, \**p*≤0.05, \*\**p*≤0.01, \*\*\**p*≤0.001, \*\*\*\**p*≤0.0001.

### *Igh* sequencing of reactivated plasmablasts suggests that heterologous priming did not significantly impact the repertoire diversity or shared clones compared to influenza infection alone

To determine if heterologous infection/immunization priming of CD4+ T cells markedly impacted the B cell clonal repertoire selection compared to mice infected with only influenza, we used the priming and influenza challenge experimental setups as previously described in Figure 2A, then 100 days after influenza challenge, mice were immunized i.p. with 10 μg recombinant HA (rHA) from PR8 influenza without adjuvant (**Fig 7A**) to preferentially engage HA-specific memory B cells to analyze secondary plasmablasts derived from the recalled HA-specific B cells. Five days after rHA immunization, IgD– CD19+B220^high/low^Fas+CD138+ plasmablasts were sorted from spleens (**Figs 7A and S6A**). Genomic DNA was isolated from sorted plasmablasts and *Igh* amplification and sequencing were performed using the immunoSEQ platform from Adaptive Biotechnologies.

**Figure 7.**
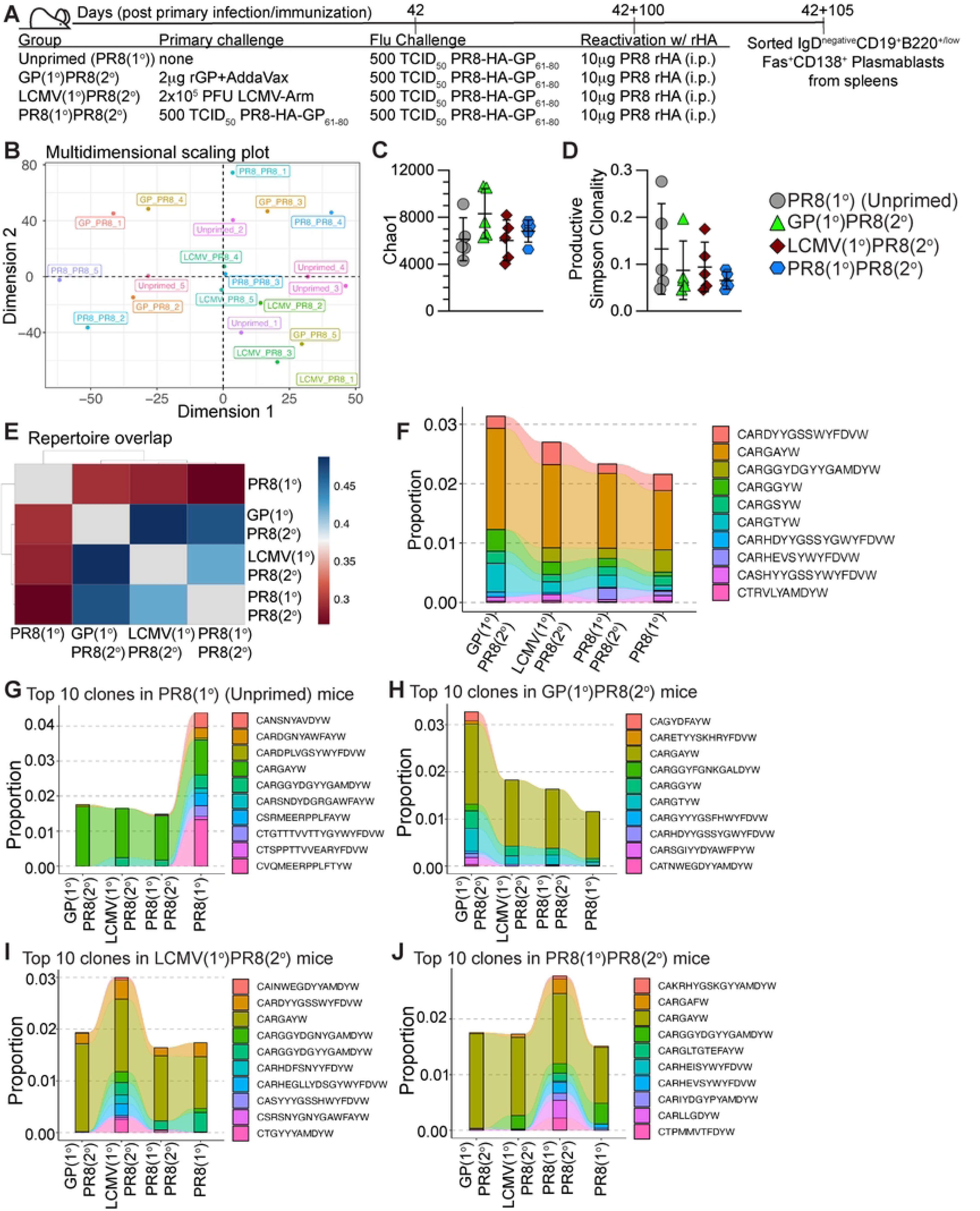
Generation of memory CD4+ T cells by heterologous immunization did not significantly impact long-lived plasmablast repertoire diversity compared to influenza infection alone. 105 days after influenza infection, unprimed mice (PR8(1°), unfilled circle), heterologously primed mice (GP(1°)PR8(2°), filled triangle, or LCMV(1°)PR8(2°), filled diamond), and homologously primed mice (PR8(1°)PR8(2°), filled hexagon) were immunized i.p. with 10 μg rHA to reactivate influenza-specific plasmablasts. 5 days postimmunization with rHA, IgD–CD19+B220+/low Fas+CD138+ plasmablasts were sorted from spleens and genomic DNA was isolated for *Igh* sequencing. **(A)** Schematic of experimental design. **(B)** Multidimensional scaling plot of CDR3 amino acid sequence repertoire overlap of individual mice. **(C)** Chao1 estimation of *Igh* repertoire diversity of plasmablasts from individual mice. **(D)** Simpson clonality diversity measure for all productive rearrangements of individual mice. **(E)** Heatmap of repertoire overlap analysis by Morisita overlap index of all mice pooled for each infection group. **(F)** Clonotype tracking analysis across priming groups of the ten largest CDR3 (amino acid sequence) clones by proportion of productive frequency shared in ≥5 infected mice (“public” clones). The productive frequency proportion value is the sum of a clone’s productive frequency in all individual mice. **(G-J)** Clonotype tracking analysis across priming groups of the ten largest CDR3 (amino acid sequence) clones by proportion of productive frequency shared in ≥2 mice in one priming group. **(G)** Ten largest CDR3 clones by proportion of productive frequency shared in ≥2 mice from the PR8(1°) (Unprimed) group. **(H)** Ten largest CDR3 clones by proportion of productive frequency shared in ≥2 mice from the GP(1°)PR8(2°) group. **(I)** Ten largest CDR3 clones by proportion of productive frequency shared in ≥2 mice from the LCMV(1°)PR8(2°) group. **(J)** Ten largest CDR3 clones by proportion of productive frequency shared in ≥2 mice from the PR8(1°)PR8(2°) group. *n* = 5 per group. Data shown are from one independent experiment. Error bars (panels C and D) are Mean±SD.

To investigate the overlap of individual mice repertoires, we performed multidimensional scaling (MDS) analysis on CDR3 amino acid sequences using the overlap coefficient of the *repOverlap* function of the Immunarch (58) package. There was no discernible clustering by priming strategy by MDS analysis (**Fig 7B**), indicating there was no significant impact on the repertoires of mice primed by the same strategy. We performed Chao1 estimation (76) and found no differences in clonal repertoire diversity richness by priming strategy (**Fig 7C**). We next analyzed the diversity of the productive rearrangements (rearrangements that produce functional B cell receptors) in individual mice by Simpson clonality measure, which is calculated as the square root of Simpson’s Index (77), which suggested that all repertoires skewed more polyclonal than mono- or oligoclonal (**Fig 7D**). When we compared the number of unique CDR3 amino acid sequence clonotypes to total clonotypes in individual mice, we found unique clones accounted for 85-95% of every individual repertoire (**Fig S6B**). In addition, we analyzed CDR3 clonotypes for differences in amino acid sequence length and number of somatic hypermutations (SHM) within nucleotide sequences and found no differences when compared by priming strategy (**Figs S6C-D**). We next performed the Morisita overlap index test (78–81) on CDR3 amino acid sequences pooled for all 5 mice in each group to evaluate the repertoire overlap by priming strategy. Our data suggest the PR8(1°) group had the most unique repertoire, while the GP(1°)PR8(2°) and LCMV(1°)PR8(2°) groups had more similar repertoires to one another (**Fig 7E**). When we analyzed the amino acid CDR3 sequences of individual mice with the Morisita overlap index test, we found that most of the GP(1°)PR8(2°) and LCMV(1°)PR8(2°) mice were more similar to each other than mice only infected with influenza (**Fig S6E**).

To evaluate shared CDR3 sequences and investigate proportional differences in mice by priming strategy, we tracked the largest 10 clonotypes by total proportion and shared in at least 5 of 20 total infected mice (“public” clonotypes) using the *trackClonotypes* feature of the Immunarch (58) package (**Fig 7F**). We found trending differences in clonotype proportions, including increased proportions of the CARGGYW and CARGTYW clones and a lack of the CARGGYDGYYGAMDYW clone in the GP(1°)PR8(2°) group (**Fig 7F**). In addition, the CARHEVSYWYFDVW clone was found in mice only infected with PR8-HA-GP_61-80_ (3 of 5 PR8(1°) mice and 4 of 5 PR8(1°)PR8(2°) mice) (**Fig 7F**). Only the CARGAYW clone was shared in all 20 infected mice, and thus had the largest representation by proportion (**Fig 7F**). Additionally, this clone was not contained in our control naïve CD19+Fas–IgD+ B cells (data not shown). We then analyzed the 10 clonotypes largest by proportion in each priming strategy group shared in at least 2 of 5 mice (**Figs 7G-J**). Our data showed that the largest shared clone, CARGAYW, was less represented proportionally in the PR8(1°) group while unique clones, including CVQMEERPPLFTYW, were more largely represented (**Fig 7G**). In addition, our data show the 10 proportionally largest clones in the PR8(1°) group comprised over 2-fold more of the total pooled repertoire proportion (>4% total) compared to the other three groups (all <2% total) (**Fig 7G**), and with the Morisita overlap data (**Fig 7E**) suggests a more unique repertoire for the PR8(1°) group. Together, these data show that while the plasmablast repertoires of individual mice were dominated by unique clones, analyses of the shared clones among individual mice were able to characterize differences in the representation of specific clonal sequences by priming strategy.

## Discussion

Current vaccine strategies, including seasonal influenza vaccines, are not specifically designed to engage CD4+ T cells, despite their necessity in germinal center formation and long-lived humoral immunity, as well as their contribution to cellular immunity in infected tissues. In this study, we used heterologous priming with adjuvanted rGP immunization or LCMV intranasal infection to generate memory CD4+ T cells and investigate the effects of recalled memory CD4+ Tfh cells and established T_RM_ cells on the response to influenza challenge. Our findings demonstrated that heterologous infection/immunization priming induced a population of antigen-specific memory CD4+CXCR5+ Tfh cells that were successfully recalled to secondary effector GC Tfh cells and induced an increased magnitude of HA-specific GC B cells compared to primary influenza infection. Furthermore, while LCMV-primed mice had significantly higher GC Tfh cells 8 dpi, our data suggested rGP-immunization priming produced higher quality GC Tfh cells as these mice had a significantly higher ratio of HA-specific GC B cells to GC Tfh cells. Heterologous infection/immunization priming also induced increased secondary effector CXCR5– Th1 cells that expressed both TBET and IFNγ, which were maintained at a higher magnitude even at memory. In addition, heterologous infection/immunization priming generated an increased long-lived CD4+ T_RM_ pool and induced increased expansion of recalled antigen-specific CD4+ T cells in the lung after influenza challenge. Interestingly, the skewing of lung-infiltrating CD4+ T cells was dependent on priming activation, as rGP immunization-primed mice preferentially recalled Tfh-like cells compared to LCMV-primed mice that preferentially recalled Th1-like cells. However, despite the early enhancement of the germinal center cellular response after influenza challenge, heterologous infection/immunization priming of CD4+ T cells did not enhance HA-specific antibody titers. Overall, our findings suggest that heterologous infection/immunization priming of CD4+ T cells can be used to enhance both the early GC response, including the GC Tfh and GC B cell magnitude, and establishment of CD4+ T_RM_ cells that respond to influenza challenge.

Tfh cells have been shown to be the limiting cell subset in the GC reaction and critical for the B cell maturation processes and production of high affinity antibodies (48–50). Our study specifically aimed to investigate whether altering the magnitude of memory CD4+ T cell help in the GC reaction would enhance the generation of antiviral humoral immunity to primary influenza infection. Previous studies have established that lineage-committed memory Th1 and Tfh cells generated during intracellular pathogenic infections can be specifically recalled upon subsequent challenges (65, 82–85). In addition, increases in Tfh cells have been shown to positively correlate with increases in GC B cell magnitude and broadly neutralizing antibodies in response to viral infections and vaccinations (40–44, 86–102). Preclinical studies investigating novel vaccination strategies successfully targeted increases in antigen-specific Tfh cells and GC and humoral responses (103–106), signifying the importance of targeting CD4+ T cells in the GC and production of high affinity antibodies. However, vaccination strategies or adjuvants specifically to target the recall of CD4+ Tfh cells to enhance the GC and its products have been slow to develop beyond the preclinical stage. We found that specifically targeting the recall and expansion of memory antigen-specific CD4+ T cells induced an increase in GC Tfh and HA-specific GC B cells early compared to mice that lacked antigen-specific memory CD4+ T cell during primary influenza infection, indicating that our findings concur with previous studies that targeting CD4+ T cells is a successful strategy to enhance the GC reaction (40, 44, 107). While heterologous infection/immunization priming enhanced the early GC Tfh and GC B cell magnitude, we did not see increases in HA-specific antibody titers, but we did see selection for specific clones in the resultant B cell repertoires. Overall, our study demonstrates that increasing the amount of antigen-specific Tfh cell help can drive an increase in the size of the germinal center response to infection, indicating that future studies could use heterologous infection/immunization priming of T helper cells to improve humoral immune responses.

Previous studies investigating T cell responses after intranasal immunization showed induction of protective proinflammatory lung-resident antigen-specific CD4+ and CD8+ T cells early after influenza challenge (69, 108–112). As virus- and vaccine-induced lung-resident CD4+ T_RM_ cells have been shown to mediate protection from influenza infection (66–70), it is important to understand how heterologous infection/immunization priming of CD4+ T_RM_ cells and resultant subsets of T_RM_ cells could enhance localized immune responses. CD4+ T_RM_, specifically resident Tfh cells, have also been shown to be important in the formation of and promotion of CD8+ T cell and B cell localization to inducible BALT structures (71, 113–115). Regarding the importance of CD4+ T cells in formation of iBALT tertiary germinal center-like structures (116–121), and that we saw increases in lung-resident Th1 and Tfh-like cells in heterologously primed mice, the differences in cellular composition or iBALT formation kinetics with either LCMV infection or adjuvanted rGP immunization priming compared to primary influenza infection warrants further investigation. As we have previously shown, non-Tfh cell populations are different between adjuvanted rGP immunization and LCMV infection (60), investigating the CD4+ T_RM_ subsets resultant from these priming strategies and their distinct roles in their recall during an influenza challenge poses an interesting question, as IFNγ-secreting CD4+ T cells have been shown to be protective against influenza infection in secondary recalled responses (122, 123).

Prior studies investigating the recall of memory CD8+ T cells in heterosubtypic influenza infection have shown that protective CD8+ T_RM_ cells were found to undergo robust clonal expansion after secondary infection and express large amounts IFNγ, though the secondary effectors were dominated by recognition of a single immunodominant epitope (124–128). As one study found neither infection of the lung nor antigen persistence was required for establishment in the lung of antigen-specific CD8+ T cells (126), we found similar results in our study investigating CD4+ T cells as adjuvanted rGP immunization showed minimal lung-resident memory CD4+ T cells prior to influenza challenge but had significantly expanded secondary effector CD4+ T cells and CD4+ T_RM_ in the lung compared to primary influenza infection, suggesting either increased trafficking to the lung or a larger antigen-specific memory T_RM_ pool compared to naïve mice.

In agreement with the idea that Tfh cells are the limiting cell subset in the GC reaction and the generation of GC-derived products, following heterologous influenza rechallenge of memory CD4+ T cells we saw an early increased magnitude of antigen-specific GC Tfh and GC B cells. However, additional studies are needed to assess the direct impact of heterologous infection/immunization priming of CD4+ T cells on survival, protection, or enhancing production of cross-reactive high affinity antibodies in response to influenza challenge. By investigating GC Tfh cell involvement in the enhancement of antiviral humoral immune responses, it is evident that new vaccination strategies should be specifically designed to engage memory CD4+ T cells to enhance the GC. Furthermore, our findings that heterologous infection/immunization priming increased expansion of localized lung antigen-specific CD4+ immune responses and lung T_RM_ populations suggest understanding differences in the lung-resident CD4+ T cell responses induced by vaccination versus previous viral infection may also be important in novel vaccine design. Ultimately, future studies are necessary to determine the mechanisms into the direct involvement of naïve versus pre-existing memory Tfh cells in preferentially generating universal and broadly neutralizing antibodies to enhance protection against influenza infection or in development of novel vaccine strategies.

## Author contributions

Conceived and designed the experiments: LMS, AGR, HJ, MAW, JSH. Performed the experiments: LMS, AGR, HJ, AB, MAW, JSH. Analyzed the data: LMS, AGR, HJ, MAW, JSH. Provided critical reagents and materials: IM, AG-S. Wrote the paper: LMS, AGR, MAW, JSH. Supervision and oversight: MAW, JSH.

## Acknowledgements

Flow cytometry data collection and cell sorting for this publication were supported by the University of Utah Flow Cytometry Core Facility. The authors thank Dr. Carl Davis at Emory University for providing the 293A-sGP cell line and glycoprotein purification protocol. The authors thank Dr. Ali Ellebedy and Dr. Jackson Turner at Washington University School of Medicine for the hemagglutinin biotinylation and staining protocol and useful discussions. The authors thank Dr. Florian Krammer at Icahn School of Medicine at Mount Sinai for the mouse-adapted PR8 and PR8-HA-GP_61-80_ viruses.

